# A single session of mild intensity physical exercise modulates brain oscillations in healthy young adults: a pilot study

**DOI:** 10.1101/2025.10.10.681596

**Authors:** Jesús Cespón, Ignacio Torres-Alemán

**Affiliations:** Department of Clinical Psychology and Psychobiology, University of Santiago de Compostela, Santiago de Compostela, Spain; Institute of Psychology (IPsiUS), University of Santiago de Compostela, Santiago de Compostela, Spain; Achucarro Basque Center for Neuroscience, Leioa, Spain

## Abstract

An acute session of moderate or vigorous physical exercise (PE) induces a cascade of neurophysiological processes such as release of growth factors, which relate to increased electroencephalogram (EEG) activity. Studies using animal models of Alzheimer’s disease (AD) showed that these mechanisms are disrupted even at asymptomatic stages of the disease. Specifically, increased neural activity within Theta band observed in healthy mice was not evidenced in mice models of AD, suggesting that EEG could be a suitable non-invasive tool to detect preclinical AD. The present study aims investigating the possible neurophysiological effects after a session of mild intensity PE, which is feasible to carry out in most population, during an EEG recording. Thus, sixteen young humans cycled at a low intensity in a stationary bike to study PE effects on the EEG frequency bands. EEG was acquired before and after PE (immediately after performing the PE, or 20-25 minutes later). Results showed that PE increased Alpha activity in frontal and central electrodes for at least 25 minutes, which aligns with previous studies in humans. Trends to increased Theta activity were observed within the left hemisphere immediately after PE, but not 25 minutes after finishing PE. Studies using larger samples should assess whether mild intensity PE increases Alpha and Theta and induces effects of different duration in both frequency bands, suggesting sensitivity of EEG to detect diverse neurophysiological effects induced by PE. Another pending issue is whether increased Alpha after PE in humans is functionally equivalent to increased Theta observed in mice.

## INTRODUCTION

The physical exercise (PE) has important neuroprotective effects (Cotman and Berchtold, 2002) and it stimulates cognition regardless the exercise is performed in a chronic or in an acute way (Basso and Suzuki, 2017; Nakajima et al., 2010; Yanagisawa et al., 2010). These beneficial properties of PE were observed in animal models (Carro et al., 2001) as well as in human beings (Lippi et al., 2020; Mandolesi et al., 2018). An acute session of PE is enough to produce a cascade of neurophysiological processes such as release of growth factors (e.g., brain-derived neurotrophic factor (BDNF), and insulin-like growth factor-1 (IGF-1)), which are related to functional changes and improved cognition after performing PE (Chieffi et al., 2017; Kandola et al., 2016; Stillman et al., 2016). These neural modulations induced by PE can be measured in human beings in a non-expensive and non-invasive manner by means of the electroencephalogram (EEG) technique (Hosang et al., 2022; Nunez et al., 2003).

Research has suggested that characterizing the PE after-effects could be useful to develop non-invasive diagnostic tools based on EEG signal because it can reflect several neurophysiological processes triggered by PE. In this context, studies about PE after-effects in animal models suggested that PE stimulates cognitive function in healthy mice but not in mice models of AD even if they are still asymptomatic (Miki Stein et al., 2017). These mentioned improvements link to specific changes in brain activity patterns, which are reflected in the EEG as increased activity within the Theta frequency band (Miki Stein et al., 2017). The neurophysiological mechanisms underlying lack of increased Theta activity in mice models of AD are thought to be a reduction in the entry of IGF-1 from blood to brain (Carro and Torres-Aleman, 2004; Trueba-Saiz et al., 2013). This statement is supported by findings about reduced correlations between IGF-I measured in blood and in the cerebrospinal fluid in mice models of AD (Trueba-Saiz et al., 2013), and it is consistent with reduced brain plasticity observed in AD patients after implementing interventional programs based on PE (Cespón et al., 2018).

There is a considerable number of studies carried out in healthy humans that investigated the effects induced by PE on neural activity by analyzing the EEG signal during a resting state (for a review, see Hosang et al., 2022) as well as examining event-related brain potentials (ERP) during the performance of a variety of cognitive tasks (for a review, see Gusatovic et al., 2022). Overall, these studies reported that an acute session of PE lead to earlier ERP latencies and increased ERP amplitudes during the performance of cognitive tasks (Gusatovic et al., 2022). Importantly, these ERP patterns (i.e., earlier ERP latencies and increased ERP amplitudes) usually correlate with enhanced cognitive performance (Cespón et al., 2023; Zhunussova et al., 2024). In addition, during the resting state, studies usually reported that PE increases neural activity within Alpha band (e.g., Bailey et al., 2008; Brümmer et al., 2011; Chaire et al., 2020; Gutmann et al., 2018; Wollseiffen et al 2016; Bailey et al., 2008; Contreras-Jordán et al., 2022), with a fair number of studies also reporting increased Beta band activity (e.g., Devilbiss et al., 2019; Contreras-Jordán et al., 2022; Guimaraes et al., 2015; Schneider et al., 2009), and some research also reporting increased Theta activity (Contreras-Jordán et al., 2022).

In sum, the results described in the previous paragraphs suggest that investigating effects induced by an acute session of PE may represent a promising avenue to distinguish between healthy aging and AD at pre-symptomatic stages of the disease. Nonetheless, investigating PE after-effects on EEG usually involves implementing a session of moderate or vigorous PE, which may be unfeasible in many participants or patients (particularly, in older adults with fragility conditions). Thus, it would be important to determine whether and to what extent effects induced by PE appear after a session of mild intensity PE, which is realistic to implement in most people. Also, investigating how long may last the effects induced by PE will provide information about the timing when such effects must be assessed. This information may be particularly relevant in studies using cognitive tasks to investigate whether an acute session of PE improves specific cognitive functions. Specifically, a task duration exceeding the time window of the PE after-effects might erroneously lead to infer the absence of cognitive improvements after an acute session of PE.

In the present study, we recruited a sample of healthy young adults. The main objective of this pilot study was investigating whether a session of mild PE modulates the EEG resting state. Although brain activity modulation by moderate and vigorous PE has repeatedly been reported, we were interested to test whether mild PE can modulate brain oscillations because a mild level of intensity would be ideal to eventually develop a test to screen the general population. According to previous studies in humans, it was expected that an acute session of mild PE would increase activity within Alpha frequency band. Additionally, we explored whether activity is also increased within Beta and Theta frequency bands because it has also been reported by several investigations. Another objective was examining, in the studied frequency bands, the duration of the possible effects induced by PE. For this purpose, EEG resting state was carried out immediately after finishing the PE in a half of the sample and 20-25 minutes after finishing the PE in the other half of the sample.

## METHODS

### Participants

Sixteen healthy young adults aged between 22 and 34 years old were recruited from the general population. Two participants were removed due to technical problems in the EEG acquisition. Consequently, the final sample was compounded by 14 healthy young adults (10 women; 4 men; mean age = 26.3; Standard deviation: 4.06). All the volunteers were free of any medical or psychiatric condition that could interfere with their participation in the study. The objectives and experimental procedures were comprehensively explained to all the participants, who provided written informed consent before the beginning of the experimental session. The study was carried out according to the principles established in the Declaration of Helsinki (1964). Moreover, the study received approval from the local ethical committee of the institution where it was implemented (Basque Center on Cognition, Brain and Language).

### Experimental procedures

Participants sat in a comfortable armchair while an EEG electrode cap was placed. Next, participants were instructed to perform the experimental session, which consisted of EEG resting state for 5 minutes and a cognitive task (speech perception task) for about 20 minutes. During the resting state, participants were requested to direct their gaze to the lower part of the computer screen to minimize ocular movements. In contrast to eyes closed resting state, eyes open resting state maximized comparability with task-related oscillations (even if it was not the focus of the present study). After finishing the resting state and the cognitive task, participants performed physical exercise (PE) in a stationary bicycle for 30 minutes at a mild intensity in such a way that they could continuing to talk during the exercise but feeling a little bit tired at the end (they filled out the Borg self-assessment physical effort questionnaire afterwards). The time duration of 30 minutes was chosen because some studies suggested that it represents a threshold to produce a substantial amount of neurotrophic factors (Lippi et al., 2020). The mild (or mild-to-moderate) exercise parameters were displaying 100-110 beats/min and a value about 10-12 on the Borg scale (a scale of perceived effort). Therefore, we have taken one objective and one subjective measure of PE intensity. During the PE, the first 3 minutes served as adaptation and the last 3 to gradually stop. After the PE, participants performed again the protocol applied before the PE. That is, participants performed the resting state and the cognitive task during the EEG recording. A medical practitioner monitored the participants during all the experimental session. The total duration of the experimental session was about 2 hours and 15 minutes (including EEG cap montage).

The order of the EEG resting state and the cognitive task was counterbalanced among the participants. Thus, the half of the sample performed the experimental session in the following order: EEG resting state – EEG task – PE – EEG resting state – EEG task (it has been labelled as counterbalancing 1 in the Figure 1) and the other half of the sample performed the experimental session in the following order: EEG task – EEG resting state – PE– EEG task – EEG resting state (it was labelled as counterbalancing 2 in the Figure 1). The structure of the experimental session (including the two types of counterbalancing) is graphically represented in the Figure 1.

**Figure 1.**
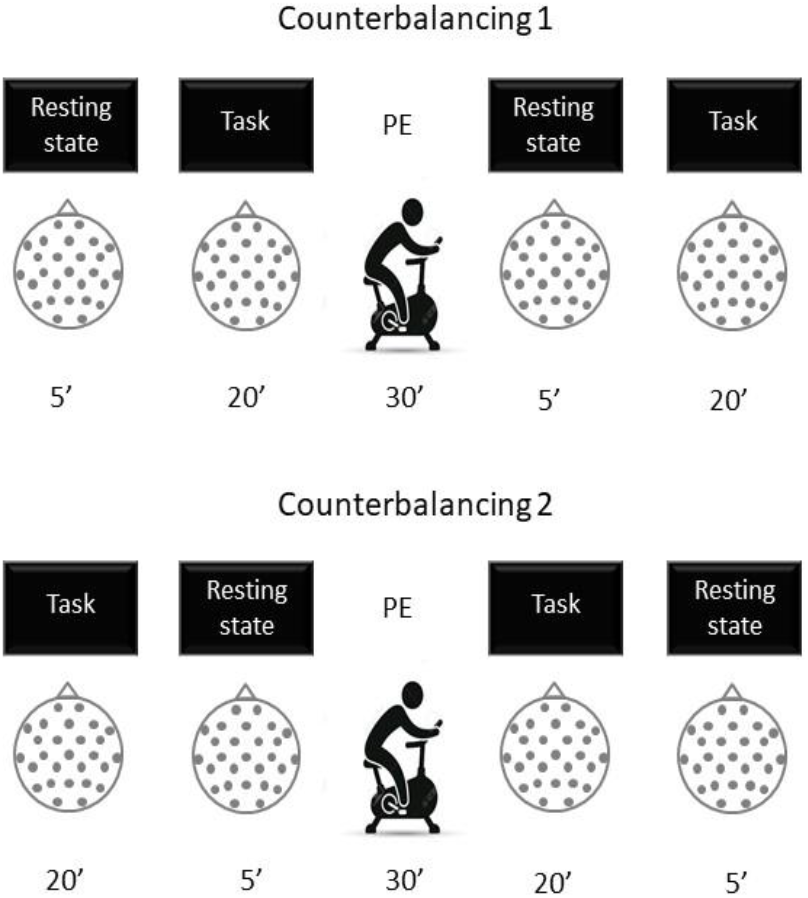
Structure of the experimental sessions. The half of the sample performed the EEG resting state immediately after doing the physical exercise (labelled as “counterbalancing 1”) whereas the other half of the sample performed the EEG resting state between 20 and 25 minutes after doing the physical exercise (labelled as “counterbalancing 2”).

### EEG recording

The EEG recording was carried out by using a 32 channels EEG cap Easycap (Brain Products GmbH, Germany). To record the EEG signal, 27 scalp electrodes were placed according to 10-20 International System of electrodes placement in the following locations: Fp1, Fp2, F3, F4, C3, C4, P3, P4, 01, O2, F7, F8, T7, T8, P7, P8, Fz, Cz, Pz, FC1, FC2, CP1, CP2, FC5,FC6, CP5, and CP6. Two vertical and two horizontal electro-oculogram (EOG) electrodes were placed in the external canthus and above-below the right eye to monitor ocular movements. Fpz was chosen as the location for the ground electrode. The left mastoid electrode was used offline to re-reference the scalp recordings to the average of the left and the right mastoid. The EEG signal was acquired using a 0.1-500 Hz bandpass filter and digitized at 1000 Hz sampling rate. Impedance was maintained below 5 kΩs.

### Data analysis

The EEG signal was filtered with a 0.1-40 Hz digital bandpass filter and a 50 Hz notch filter. Horizontal and vertical EEG ocular artifacts as well as muscular artifacts were removed from the EEG signal by using Independent Component Analysis (ICA). Subsequently, the EEG was segmented in epochs of 2 seconds, giving rise to 150 segments. Segments containing artifacts were removed by automatically rejecting any activity exceeding 50μV. All remaining epochs were visually inspected and removed from further analysis is still displayed any artifact. Next, Fourier transform was used to obtain the spectrum of frequencies (Delta, Theta, Alpha, and Beta). Spectral activity was calculated in the 27 electrodes placed on the scalp for the two bands of interest; that is, Theta (4.1-7.9Hz), Alpha (8.6-11.9Hz), and Beta (12.9-24.6Hz). These analyses were run by using the software Brain Vision Analyzer 2.3 (Brain Products, GmBh, Gilching, Germany).

### Statistical analysis

Paired sample t-tests were carried out for Alpha, Theta, and Beta activity to assess whether activity changed after PE in several representative scalp locations (specifically: F3, Fz, F4, C3, Cz, C4, P3, Pz, and P4). Moreover, independent sample t-tests were carried out to compare the PE-related effects in those subjects who conducted the resting state immediately after the PE (counterbalancing 1 or group 1) and those subjects who carried out the resting state 20-25 minutes after the PE (counterbalancing 2 or group 2). For this purpose, we subtracted, for each participant and analyzed frequency band, activity measured after doing the PE from the activity measured before performing the PE.

With the aim of deepening on possible group-related differences between participants who performed the resting state immediately after the PE and participants who performed the resting state 20-25 minutes after the PE, repeated measures ANOVAs were carried out, for Alpha, Beta, and Theta activity, using an inter-subject factor: Counterbalancing, or Group (two levels: Group 1, Group 2) and the following intra-subject factors: Time (two levels: before PE, after PE), and Electrode (three levels: F3, F4, C3). The electrodes were selected ad hoc for being representative electrodes that displayed increased activity after PE, as revealed by paired sample t-tests. When ANOVAs revealed significant effects, pairwise comparisons were performed using the Bonferroni correction. Also, when sphericity condition was not met, Greenhouse-Geiser correction was applied for the degrees of freedom. Partial eta squared (η^2^p) was used to estimate the effect size. All statistical analyses were carried out using SPSS statistical software version 18 (IBM Corporation, Armonk, NY, USA). The statistical significance was determined by using an alpha level of 0.05.

## RESULTS

### t-tests

Paired sample t-tests for activity within Alpha frequency band showed significant effects in the following electrodes: F3: t(13) = 3.469, p = 0.004; F4: t(13) = 3.121, p = 0.008; C3: t(13) = 2.786, p = 0.015; Fz: t(13) = 3.232, p = 0.007. These results, represented in the Figure 2, indicated increased Alpha activity after PE. Paired sample t-tests for activity within Theta frequency band revealed marginally significant effects in the following electrodes: F3: t(13) = 1.931, p = 0.076; C3: t(13) = 2.077, p = 0.058; P3: t(13) = 2.103, p = 0.055. These results, which are graphically represented in the Figure 2, indicate a trend to enhanced Theta activity after performing the session of PE. Paired sample t-tests for activity within Beta frequency band did not reveal any significant effect in the analyzed electrodes.

**Figure 2.**
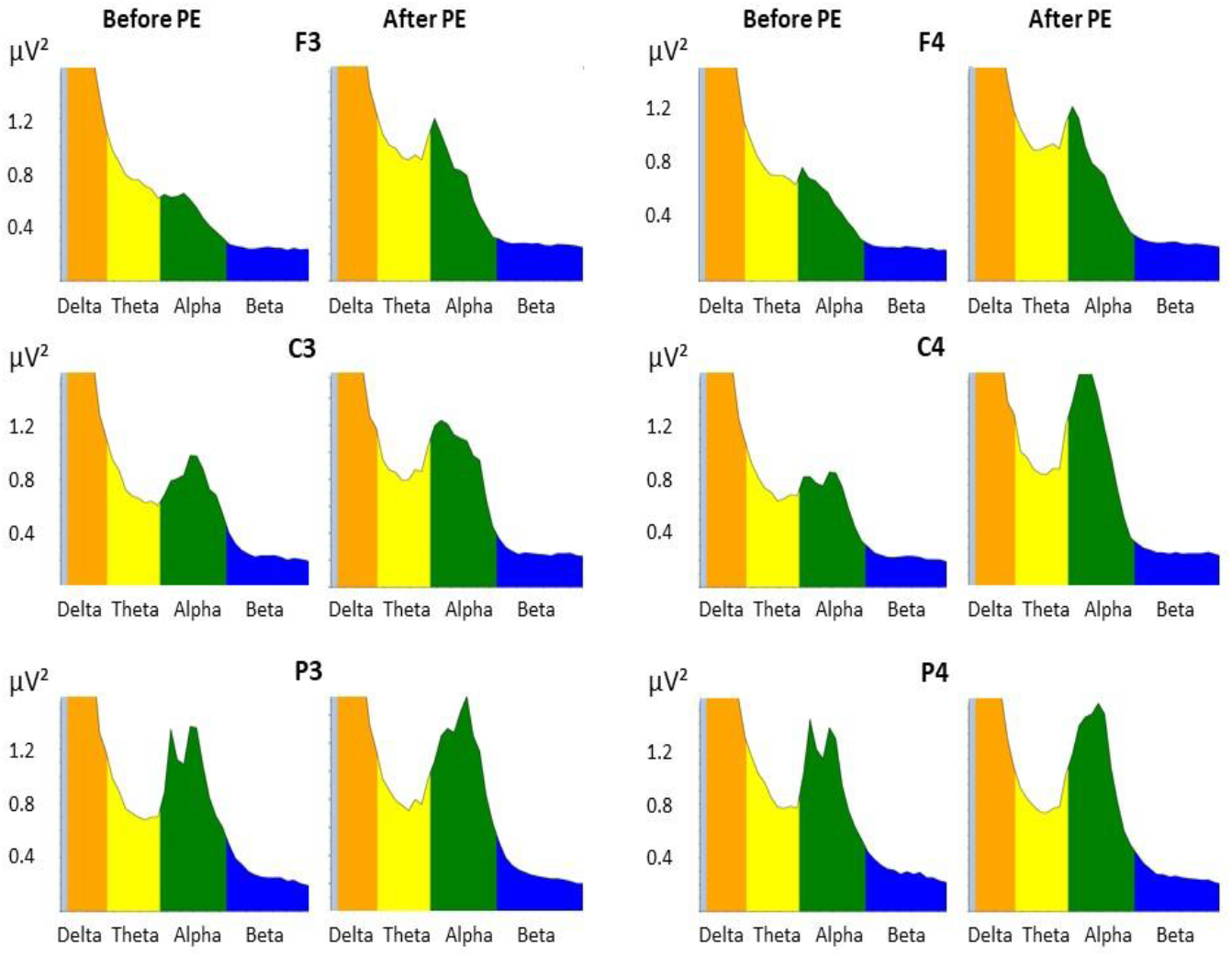
EEG resting state at representative electrode sites. This figure represents the main spectral frequency bands (i.e., delta, theta, alpha, and beta) in several representative scalp positions (namely, electrodes F3, F4, C3, C4, P3, and P4). Hypotheses and analyses focused on Beta, Alpha, and Theta frequency bands. Alpha activity was significantly increased after PE. There was also some evidence for increased Theta activity after performing the physical exercise. Beta activity changes related to PE were not statistically significant.

We used independent sample t-tests to compare the increase of Theta activity between participants performing the EEG resting state immediately after PE (counterbalancing 1) and participants performing the EEG resting state 20-25 minutes after PE (counterbalancing 2). The electrodes of interest were F3, C3, and P3. The increase of neural activity within Theta frequency band after PE was larger in counterbalancing 1 compared to counterbalancing 2 in F3 [t(12) = 2.365, p = 0.036)] and C3 [(t(12) = 2.206, p = 0.048)]. These results are graphically represented in the top charts of the Figure 3 (counterbalancing 1 on the left panel and counterbalancing 2 on the right panel).

**Figure 3.**
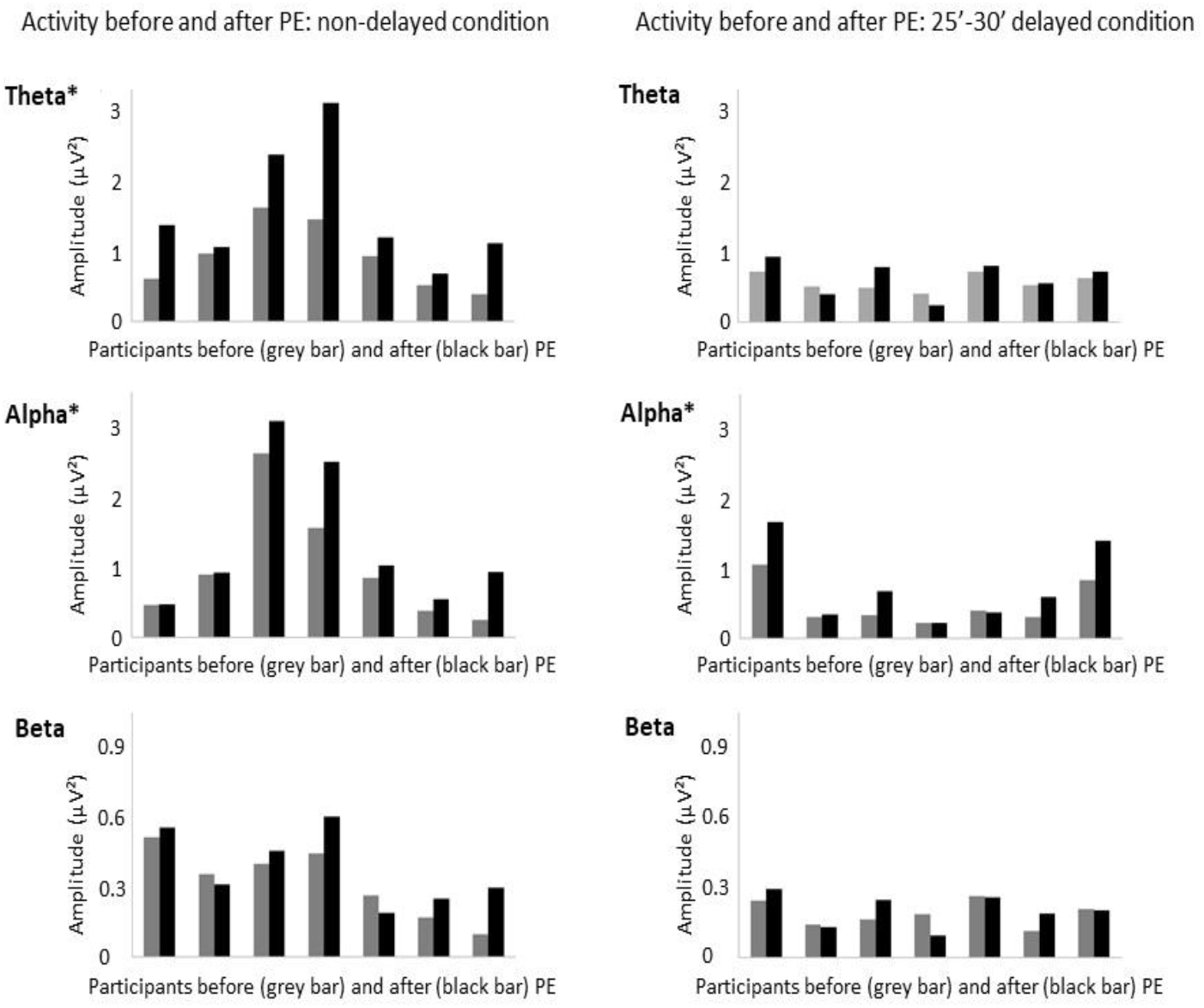
Resting state activity before and after physical exercise (PE) for each participant. This figure represents, for each participant, Theta, Alpha, and Beta activity before and after performing the PE. The representation is the result of averaging activity across F3, Fz, and F4 electrodes. This Figure distinguishes participants who performed the counterbalancing 1 (i.e., participants in the Group 1, who performed the EEG resting state immediately after doing the PE) and counterbalancing 2 (i.e., participants in the Group 2, who performed the EEG resting state about 25 minutes after doing the PE). The asterisk (*) represents that differences between activity before and after PE were statistically significant.

Independent sample t-tests carried out in F3, F4, Fz, and C3 electrodes (i.e., where paired t-tests revealed increased neural activity after PE in Alpha band) did not reveal significant differences in the increase of neural activity within Alpha and Beta frequency bands regardless the EEG resting state was recorded immediately after PE (i.e., in the counterbalancing 1) or 25 minutes later (i.e., in the counterbalancing 2). The Figure 3 represents these results for Alpha (middle charts) and Beta (bottom charts) activity.

### Analysis of variance

The repeated measures ANOVA (Group x Time x Electrode) for Theta activity revealed an effect of Time in the limit of the statistical significance [F (1, 12) = 4.69, p = 0.05, η^2^p = 0.281], as Theta activity was higher after than before PE (p = 0.05). Moreover, the analysis revealed an effect of Group (i.e., counterbalancing) [F (1, 12) = 8.40, p = 0.013, η^2^p = 0.412] because Theta activity was higher in the group of participants who performed the EEG resting state immediately after the PE compared to group of participants performing the PE about 25 minutes after finishing the PE (p = 0.013). A marginal Time x Group interaction effect was observed [F (1, 12) = 4.12, p = 0.064, η^2^p = 0.257]. This interaction effect was driven by increased Theta activity after PE in the Group 1 (p = 0.012) but not in the Group 2.

The repeated measures ANOVA (Group x Time x Electrode) for Alpha activity revealed a significant effect of Time [F (1, 12) = 17.26, p = 0.001, η^2^p = 0.590], because neural activity within the Alpha frequency band was larger after performing the PE (p = 0.001). The analysis did not reveal other effects of Time or Group or their interactions. The repeated measures ANOVA (Group x Time x Electrode) for Beta activity did not reveal a significant effect of Time or their interactions.

## DISCUSSION

The results of the present study evidenced that a mild session of PE increased EEG resting state activity in the Alpha frequency band. Non-significant trends to increased Theta activity were also observed, whereas no differences were found in Beta band. For Alpha band, resting state activity mainly increased in bilateral frontal regions. For Theta band, resting state activity increased in frontal, central, and parietal sites within the left hemisphere. Regarding the duration of these PE after-effects, no differences between the Group 1 (i.e., when the resting state was recorded immediately after the PE) and the Group 2 (i.e., when the resting state was recorded 20-25 minutes after finishing the PE) were observed after PE in Alpha band. However, Theta activity after PE was larger in the Group 1 compared to Group 2, suggesting a short duration of the possible Theta effects induced by PE.

As previously stated, Alpha activity increased after performing a bout of PE even if the intensity of the exercise was low (or low-to-moderate), which is consistent with and extend findings from previous studies usually using moderate or vigorous intensities (Hosang et al., 2022). Thus, the results of the present study suggest that a mild intensity bout of PE is enough to induce neurophysiological effects. This finding may have important clinical implications in case of developing PE tests to screen brain health because, unlike moderate or vigorous PE, a session of mild or mild-to-moderate intensity PE would be feasible to implement in most people, including older adults.

A trend to increased activity after PE was observed in the Theta frequency band. There were topographical differences for Alpha and Theta effects. Specifically, Alpha increased in bilateral frontal sites, which was consistent with a substantial number of previous studies reporting increased Alpha activity within frontal sites after PE (e.g., Brümmer et al. 2011; Hicks et al. 2018; Kao et al., 2021; Petruzzello and Landers, 1994; Schneider et al. 2009b). On the other hand, Theta activity increased in the left frontal, central, and parietal regions. Increased Theta activity after PE was recently reported (Contreras-Jordán et al., 2022) even if it represents a finding scarcely shown in previous literature (Hosang et al., 2022). It is possible that some differences between the EEG resting state acquisition protocol of the present study and protocols used in other studies (for instance, use of open eyes instead of the more common closed eyes EEG resting state) explain the disparity of results. Finally, no evidence was observed for Beta activity changes, a finding that was often reported even if using higher intensity PE protocols (Hosang et al., 2022).

Increased Alpha activity was observed for those participants performing the resting state immediately after the PE as well as for those participants performing the resting state at 20-25 minutes after finishing the PE. In contrast, a trend to increased Theta activity was observed in those participants performing the EEG resting state immediately after the bout of PE, but not at 20-25 minutes later. Future research using larger samples will be crucial to determine if these results and differences in the duration of the PE after-effects observed in Alpha and Theta bands can be confirmed. Importantly, a duration of (at least) 25 minutes for increased Alpha activity establishes a time window that could be reliably used to investigate cognitive improvements after PE linked to increased Alpha activity.

It is worthy to emphasize that, unlike the mentioned findings obtained in mice, Theta activity after PE in human young adults has barely been reported (Hosang et al., 2022). There are (at least) two possible explanations for these inconsistent findings. Firstly, there are obvious differences between mice and human brains. In fact, it has been stated that EEG rhythms are faster in humans compared to rodents for some comparable neurophysiological processes (Jacobs, 2013; Maheshwari, 2020). Thus, it is possible that increased Theta activity after PE observed in mice is functionally equivalent to increased Alpha activity after PE observed in humans. Secondly, it is possible that processes related to IGF-1 are reflected in the EEG but, at the same time, these processes are overlapped and/or partially masked by other neurophysiological processes induced by PE. For instance, early EEG studies had already reported that a bout of PE produces a release of endorphins for up to 1 hour (Sforzo, 1989), which increases activity within Alpha frequency band (Pfefferbaum et al., 1979). A possible future direction of studies using animal and human participants may be determining to what extent increased Alpha activity after PE in humans may be functionally equivalent to increased Theta activity observed in mice after PE.

The present pilot study has several limitations that must be considered. An obvious limitation of this study is the reduced sample size. Also, an optimal research approach would involve using a within-subjects design and analyzing an entire time window of at least 30 minutes to investigate the duration of the possible effects induced by a bout of mild intensity PE. Also, analyzing EEG resting state during 30 minutes following the PE can prevent effects linked to the performance of the cognitive task (Kavcic et al., 2022; Lor et al., 2023), which might have been present in the current study (note that, even if not significant, the Figure 3 suggests that baseline activity is lower in the Group 2 compared to the Group 1, which may be related to the preceding task performed by participants in the Group 2).

In summary, the results of the present study showed that a mild intensity bout of PE increases Alpha activity within bilateral frontal regions, suggesting that mild intensity PE is enough to produce neurophysiological changes in the brain. Moreover, the increase of activity in the Alpha band remained for, at least, 25 minutes after finishing the PE. It suggests a time window of (at least) 25 minutes to study the existence of cognitive improvements after a session of acute PE. In detail, a person performing any cognitive task within a time window of 25 minutes after finishing the PE would be prone to exhibit cognitive improvements linked to increased Alpha activity. Finally, the present study showed that Theta activity increased after PE, but such increment was only marginal, and it was not observed in participants performing the resting state 25 minutes after finishing the PE. Future research using larger samples may assess the strength and reliability of these findings, which suggest that mild intensity PE may trigger diverse neurophysiological processes of different duration that can be detected by using the EEG.

## ACKNOWLEDGEMENTS

This project has received funding from the Basque Government IKUR strategy (SoMenHe), and Ramón y Cajal program (JC: No. RYC2022-035443-I). The funding sources were not involved in any activity contributing to this publication.

## REFERENCES

Bailey, S.P., Hall, E.E., Folger, S.E., & Miller, P.C. (2008). Changes in EEG during graded exercise on a recumbent cycle ergometer. J. Sports Sci Med, 7(4), 505–511. https://www.ncbi.nlm.nih.gov/pmc/articles/PMC3761919/.

Basso, J.C., Suzuki, W.A. (2017). The Effects of Acute Exercise on Mood, Cognition, Neurophysiology, and Neurochemical Pathways: A Review. Brain Plast, 28;2(2), 127–152. doi: 10.3233/BPL-160040.

Brümmer, V., Schneider, S., Abel, T., Vogt, T., & Strüder, H.K. (2011). Brain cortical activity is influenced by exercise mode and intensity. Med. Sci. Sporrts Exerc., 43(10), 1863–1872. 10.1249/MSS.0b013e3182172a6f.

Carro, E., & Torres-Aleman, I. (2004). The role of insulin and insulin-like growth factor I in the molecular and cellular mechanisms underlying the pathology of Alzheimer’s disease. Eur J Pharmacol 490,127–133.

Carro, E., Trejo, J.L., Busiguina, S., & Torres-Aleman, I. (2001) Circulating insulin-like growth factor I mediates the protective effects of physical exercise against brain insults of different etiology and anatomy. J Neurosci 25678–5684.

Carro, E., Trejo, J.L., Gomez-Isla, T., LeRoith, D., & Torres-Aleman, I. (2002) Serum insulin-like growth factor I regulates brain amyloid-beta levels. Nat Med 81390–1397.

Cespón, J., Chupina, I., & Carreiras, M. (2023). Cognitive reserve counteracts typical neural activity changes related to ageing. Neuropsychologia, 9;188:108625. 10.1016/j.neuropsychologia.2023.108625.

Cespón, J., Miniussi, C., & Pellicciari, M.C. (2018). Interventional programmes to improve cognition during healthy and pathological ageing: Cortical modulations and evidence for brain plasticity. Ageing Res Rev. 43, 81–98. doi: 10.1016/j.arr.2018.03.001.

Chaire, A., Becke, A., & Düzel, E. (2020). Effects of physical exercise on working memory and attention-related neural oscillations. Front. Neurosci. 14, 239. 10.3389/fnins.2020.00239.

Chieffi, S., Messina, G., Villano, I., Messina, A., Valenzano, A., Moscatelli, F., Salerno, M., Sullo, A., Avola, R., Monda, V., Cibelli, G., & Monda, M. (2017). Neuroprotective effects of physical activity: evidence from human and animal studies. Front. Neurol. 8, 188.

Contreras-Jordán, O.R., Sánchez-Reolid, R., Infantes-Paniagua, Á, Fernández-Caballero, A., & González-Fernández, F.T. (2022). Physical exercise effects on university students’ attention: An EEG analysis approach. Electronics, 11(5), 770. 10.3390/electronics11050770.

Cotman, C.W., & Berchtold, N.C. (2002) Exercise: a behavioral intervention to enhance brain health and plasticity. Trends Neurosci. 25, 295–301.

Devilbiss, D.M., Etnoyer-Slaski, J.L., Dunn, E., Dussourd, C.R., Kothare, M.V., Martino, S.J., & Simon, A.J. (2019). Effects of exercise on EEG activity and standard tools used to assess concussion. Journal of Healthcare Engineering, 2019, Article 4794637. 10.1155/2019/4794637.

Guimarães, T., Macedo da Costa, B., Silva Cerqueira, L., de Carlo Andrade Serdeiro, A., Augusto Monteiro Saboia Pompeu, F., Sales de Moraes, H., Meireles dos Santos, T., & Camaz Deslandes, A. (2015). Acute effect of different patterns of exercise on mood, anxiety and cortical activity. Arch. Neurosci. 2(1), e18780. 10.5812/archneurosci.18781.

Gusatovic, J., Gramkow, M.H., Hasselbalch, S.G., & Frederiksen, K.S. (2022). Effects of aerobic exercise on event-related potentials related to cognitive performance: a systematic review. PeerJ., 10, e13604. doi: 10.7717/peerj.13604.

Gutmann, B., Huelsduenker, T., Mierau, J., Strüder, H.K., & Mierau, A. (2018). Exercise-induced changes in EEG alpha power depend on frequency band definition mode. Neuroscience Letters, 662(1), 271–275.10.1016/j.neulet.2017.10.033.

Hicks, R.A., Hall, P.A., Staines, W.R., & McIlroy, W.E. (2018). Frontal alpha asymmetry and aerobic exercise: Are changes due to cardiovascular demand or bilateral rhythmic movement? Biol. Psychol. 132, 9–16. 10.1016/j.biopsycho.2017.10.011.

Hosang, L., Mouchlianitis, E., Guérin, S.M.R., & Karageorghis, C.I. (2022). Effects of exercise on electroencephalography-recorded neural oscillations: a systematic review. Int Rev. Sport Exerc. Psychol., 17(2), 926–979. 10.1080/1750984X.2022.2103841.

Jacobs, J. (2013). Hippocampal theta oscillations are slower in humans than in rodents: implications for models of spatial navigation and memory. Philos Trans R Soc Lond B Biol Sci. 369(1635), 20130304. doi: 10.1098/rstb.2013.0304.

Kandola, A., Hendrikse, J., Lucassen, P.J., & Yücel, M. (2016). Aerobic exercise as a tool to improve hippocampal plasticity and function in humans: practical implications for mental health treatment. Front. Hum. Neurosci. 10, 373.

Karran, E., & Hardy, J. (2014) Antiamyloid Therapy for Alzheimer’s disease: are we on the right road? N. Engl. J. Med. 370, 377–378.

Kavcic, V., Daugherty, A.M., & Giordani, B. (2021). Post-task modulation of resting state EEG differentiates MCI patients from controls. Alzheimers Dement (Amst). 13(1), e12153. 10.1002/dad2.12153.

Kao, S.C., Wang, C.H., Kamijo, K., Khan, N., & Hillman, C. (2021). Acute effects of highly intense interval and moderate continuous exercise on the modulation of neural oscillation during working memory. Int. J. Psychophysiol. 160(1), 10–17. 10.1016/j.ijpsycho.2020.12.003.

Lippi, G., Mattiuzzi, C., & Sanchis-Gomar, F. (2020). Updated overview on interplay between physical exercise, neurotrophins, and cognitive function in humans. J Sport Health Sci. 2020 Jan;9(1)74–81. 10.1016/j.jshs.2019.07.012.

Lor, C.S., Zhang, M., Karner, A., Steyrl, D., Sladky, R., Scharnowski, F., & Haugg, A. (2023). Pre-and post-task resting-state differs in clinical populations. Neuroimage Clin. 37, 103345. 10.1016/j.nicl.2023.103345.

Maheshwari, A. (2020). Rodent EEG: Expanding the Spectrum of Analysis. Epilepsy Curr. 20(3), 149–153. doi: 10.1177/1535759720921377.

Mandolesi, L., Polverino, A., Montuori, S., Foti, F., Ferraioli, G., Sorrentino, P., & Sorrentino, G. (2018). Effects of physical exercise on cognitive functioning and wellbeing: biological and psychological benefits. Front. Psychol. 9:50 10.3389/fpsyg.2018.00509.

Miki Stein, A., Munive, V., Fernandez, A.M., Nunez, A., & Torres Aleman, I. (2017) Acute exercise does not modify brain activity and memory performance in APP/PS1 mice. PLoS One 12e0178247.

Nakajima, S., Ohsawa, I., Ohta, S., Ohno, M., & Mikami, T. (2010) Regular voluntary exercise cures stress-induced impairment of cognitive function and cell proliferation accompanied by increases in cerebral IGF-1 and GST activity in mice. Behav Brain Res 211, 178–184.

Nunez, A., Carro, E., & Torres-Aleman, I. (2003) Insulin-like growth factor I modifies electrophysiological properties of rat brain stem neurons. J Neurophysiol 89, 3008–3017.

Petruzzello, S.J., & Landers, D.M. (1994). State anxiety reduction and exercise: Does hemispheric activation reflect such changes? Med. Sci. Sports Exerc., 26(8), 1028–1035. 10.1249/00005768-199408000-00015.

Pfefferbaum, A., Berger, P.A., Elliott, G.R., Tinklenberg, J.R., Kopell, B.S., Barchas, J.D., & Li, C.H. (1979). Human EEG response to beta-endorphin. Psychiatry Res. (1), 83–88. doi: 10.1016/0165-1781(79)90031-3.

Robertson, I.H. (2014). Right hemisphere role in cognitive reserve. Neurobiol Aging. 35(6), 1375–1385. 10.1016/j.neurobiolaging.2013.11.028.

Schneider, S., Brümmer, V., Abel, T., Askew, C.D., & Strüder, H.K. (2009). Changes in brain cortical activity measured by EEG are related to individual exercise preferences. Phsiol. Behav., 98 (4), 447–452. 10.1016/j.physbeh.2009.07.010.

Sforzo, G.A. (1989). Opioids and exercise. An update. Sports Med. 7(2), 109–124. 10.2165/00007256-198907020-00003.

Shah, H., Albanese, E., Duggan, C., Rudan, I., Langa, K.M., Carrillo, M.C., Chan, K.Y., Joanette, Y., Prince, M., Rossor, M., Saxena, S., Snyder, H.M., Sperling, R., Varghese, M., Wang, H., Wortmann, M., & Dua, T. (2016). Research priorities to reduce the global burden of dementia by 2025. Lancet Neurol 15, 1285–1294.

Stein, A.M., Silva, T.M.V., Coelho, F.G.M., Arantes, F.J., Costa, J.L.R., Teodoro, E., & Santos-Galduróz, R.F. (2018). Physical exercise, IGF-1 and cognition A systematic review of experimental studies in the elderly. Dement Neuropsychol. 2(2), 114–122. 10.1590/1980-57642018dn12-020003.

Stillman, C.M., Cohen, J., Lehman, M.E., & Erickson, K.I. (2016). Mediators of physical activity on neurocognitive function: a review at multiple levels of analysis. Front. Hum. Neurosci. 10, 626.

Trejo, J.L., Piriz, J., Llorens-Martin, M.V., Fernandez, A.M., Bolos, M., LeRoith, D., Nunez, A., & Torres-Aleman, I. (2007). Central actions of liver-derived insulin-like growth factor I underlying its pro-cognitive effects. Mol Psychiatry, 12, 1118–1128.

Trueba-Sáiz, A., Cavada, C., Fernandez, A.M., Leon, T., González, D.A., Fortea Ormaechea, J., Lleó, A., Del Ser, T., Nuñez, A., & Torres-Aleman, I. (2013). Loss of serum IGF-I input to the brain as an early biomarker of disease onset in Alzheimer mice. Transl Psychiatry. 3(12), e330. 10.1038/tp.2013.102.

Wollseiffen, P., Schneider, S., Martin, L. A., Kerhervé, H. A., Klein, T., & Solomon, C. (2016). The effect of 6 h of running on brain activity, mood, and cognitive performance. Exp. Brain Res, 234(7), 1829–1836. 10.1007/s00221-016-4587-7.

Yanagisawa, H., Dan, I., Tsuzuki, D., Kato, M., Okamoto, M., Kyutoku, Y., & Soya, H. (2010) Acute moderate exercise elicits increased dorsolateral prefrontal activation and improves cognitive performance with Stroop test. Neuroimage 50, 1702–1710.

Zhunussova, A., Pellicciari, M.C., Carreiras, M., & Cespón, J. (2024). Electrophysiological correlates of cognitive reserve in healthy older adults at different cognitive states. bioRxiv. 10.1101/2024.12.30.627948.

